# Effect of plant tissues on DNA quantity and quality of barley (*Hordeum vulgare*) validating through PCR technique

**DOI:** 10.1101/2021.10.28.466350

**Authors:** Nigussie Kefelegn, Gizachew Haile, Hemalatha Palanivel

## Abstract

The use of molecular techniques to deal with plant molecular breeding requires the extraction of genomic DNA in good quantity and quality, which can be influenced by method of extraction and source of DNA (plant species, plant part or tissues). However, this research focuses on plant tissue source and tried to describe the quality and quantity of DNA isolated from different tissues’ of barley crop. CTAB protocol was used for the isolation of DNA and both the quality and quantity of this DNA was validated through gel electrophoresis, Qubit quantification and PCR amplifications. The result showed that DNA could be successfully extracted from all tissues of plants and the yield of DNA obtained was variable ranging from 179ng μl^-1^ in stem to 750ng μl^-1^ in young leave. Band intensity of genomic and PCR amplified DNA was good for DNA isolated from young and matured leave. Faint band was observed in the PCR amplification for DNA isolated from stem but no to unreliable amplification was obtained for seeds and roots, respectively. Thus, for any molecular technique in barley crop research, the best tissue for DNA isolation using modified CTAB, is young or matured leave and alternatively stem can be used as DNA source.

## 1. Introduction

The isolation of DNA in good quantity and quality is a critical step for many procedures used in molecular genetics and in applied plant breeding (Boiteux *et al*., 1999) and also important to answer different ecological and evolutionary questions (Haig, 1998; Allen *et al*., 2006). The large number of samples often required in breeding programs that provide high quality DNA, by using rapid, simple, and inexpensive protocols (Weising *et al*., 1995). Therefore, customizing protocols are important to improve DNA isolation in order to optimize its quality and quantity for the intended objectives or downstream processing. The most common plant tissue used for DNA extraction is leave which supposed to be high mass of dividing cell and hence with much DNA to extract. However, the DNA contaminants have varied along with plant tissue type and even along with the different species (Lucas *et al*., 2019). Many studies are carried out for plant DNA extraction either to optimize the DNA extraction protocol or to select best plant tissue for quality DNA extraction (Hasan *et al*., 2012; Marsal *et al*., 2013). But optimizing DNA extraction protocol is not yet addressed in many crops as DNA quality differ among the different protocols, plant species and also plant tissue types.

### 1.1. Background

The use of molecular techniques to deal with molecular plant breeding, ecology and evolutionary process requires the extraction of genomic DNA in good quantity and quality which can be influenced by plant species and type of plant part or tissue type (Varma *et al*., 2007; Heikrujam *et al*., 2020). Younger tissues, especially the leaves, produce DNA of good quality and quantity due to large numbers of cells and small amounts of secondary metabolites (Williams *et al*. 1994). Sahu *et al*. (2012) also reported that fresh and young leaf of Mangroves gives high quality of DNA. Varma *et al*. (2007) reported that young, healthy and tender tissues, especially partially expanded leaves make an ideal choice as they can yield good quality and quantity of DNA due to a larger number of cells and less deposition of starch and secondary metabolites. In contrast, DNA extracted from mature leave has poor quality and low yield due to the presence of high concentrations of polyphenols, tannins, polysaccharides and other secondary metabolites (Elger *et al*., 2009; Moreira and Oliveira, 2011). Research done by Sahu *et al*. (2012) indicates that obtaining good quality DNA is difficult from matured leave. However, Marsal *et al*. (2013) and Marsal *et al*. (2011) reported that matured leave were source of good quality and yield of DNA suitable for PCR amplification.

Stem and root tissues are more lignified and may present secondary compounds, which can hinder the isolation of the genetic material (Lewinsohn *et al*. 1994) or inhibit the polymerase chain reaction (PCR) used to amplify the loci of interest (Schori *et al*. 2013). Some species with exposed roots have high amounts of secondary compounds that act as natural defense pesticides, mainly against herbivores. However, in palm tree, reported by Ulloa (2020), root tissue is an ideal candidate to obtain high concentrations of nucleic acids. Especially the tip of the root contains high concentration of nucleic acid. Stem can be also a good source of DNA that can be suitable for PCR amplification (Aydin *et al*., 2020). On the other hand, many crop seeds contain polysaccharides, polyphenols, mucilage, lipids, pigments many of which cause DNA extraction from seeds difficult and sometimes a research limiting step. Despite of this limitation, DNA isolation directly from seeds provides some advantages including less-time and effort, particularly for large experiments. Indeed better quality and quantity DNA extraction has been possible yet using Cetyl tri-methyl ammonium bromide (CTAB) protocol (Boiteux *et al*., 1999; Hasan *et al*., 2012; Marsal *et al*., 2013). Elsalam *et al*. (2011) was able to extracte high molecular DNA (100 to 200 ng per 100 mg) using the standard CTAB protocol from germinated wheat seeds with minimal contamination of polysaccharides and polyphenol compounds. On the other hand, breaking the cell wall with NaOH and incubation of the seed at 50°C (water bath) was described by Post *et al*. (2003) as the best method to extract DNA from barley seed.

As previously mentioned, plant tissue was the most important source of variation for DNA yield as well as quality. This can be explained by either some intrinsic tissue-specific variation (e.g., number and size of cells; ratio of mitotic to interphase nuclei; growth stage or by a particular differences in the structure or biochemical composition (polysaccharides, polyphenols and other secondary metabolites) of the tissue. DNA quality and quantity also affected by plant species diversity (Heikrujam *et al*., 2020). Different plant species have varying levels of polysaccharides, polyphenols, and other secondary metabolites (Heikrujam *et al*., 2020; Aboul-Maaty and Oraby, 2019). According to Karaca and Ince (2018) the yield of genomic DNA of onion, squash, and eggplant, pepper, tomato and cotton seed samples ranged from 61.8 μg/g in squash to 365.6 μg/g in cotton.

Quality and quantity of DNA extraction also affected by the type of methods and their efficiency can be affected by the type of plant tissue. CTAB was very effective to generated DNA from flower and seed tissue of carrot for better amplification while the number of amplicon is lower for root and leave (Boiteux *et al*., 1999). Sahu *et al*. (2012) reported CTAB (Cetyl tri-methyl ammonium bromide) protocol enables the isolation of high quality genomic DNA amenable to RAPD (Random amplified Polymorphic DNA), restriction digestion, and amplification with reduced cost and health concerns. The CTAB method described by Saghai-Maroof *et al*. (1984) gave better DNA yield in terms of quality and quantity from the study plants. Addis Ababa Science and Technology University (AASTU) molecular laboratory uses the Modified Doyle & Doyle (1987) CTAB DNA extraction method as described here in the material methods. Most of the PhD students’ uses this protocol and they have commonly extract DNA from plant leaves. This protocol was used in this study for DNA extraction from the different tissues of barley crops and both the quality and yield of DNA is determined.

### 1. 2. Statement of the Problems

Many researches have been conducted to optimizing DNA extraction techniques from different organism. They have showed, type of species, plant tissues and the methods employed in the DNA extraction can affects the quality and quantity of DNA. However, except on some horticultural crops and plant trees, detailed study has not been carried out on field crops of their part/tissues as DNA source and its consequences on the quality and quantity of DNA extracted with the different DNA extraction protocols. With this regards this research focus on one of the field crop barley and the effect of its tissue parts as a source, on DNA quality and quantity extracted using CTAB protocol.

## 2. Study Objectives

✓ Identify the best tissue type or part for best yield and quality of DNA extraction
✓ And describe the efficiency of the modified CTAB protocol

## 3. Methodology

### 3.1. Study setting

Barley crops was grown in a pot at Addis Ababa Science and Technology University (AASTU) Ethiopia. DNA was extracted from the different tissues of barley crops (roots, stems, matured leave, young leave and seeds) and the extraction process was done in three replication. For extraction, a total of 300mg of sample was taken from each tissue and equally divided in to three tubes after it is grinded. Materials and reagents recruited in extraction process are mentioned under annexes. The overall activities were carried out at AASTU molecular laboratory.

### 3.2. Laboratory testing procedures

DNA was isolated from the different tissues/parts of barley crop using modified Doyle & Doyle (1987) Cetyl tri-methyl ammonium bromide (CTAB) protocol. Matured and young leave, stems, roots and seeds were used for the isolation of DNA based on the protocol (see description below). In general, the extraction of total genomic DNA comprised of three distinct steps.

#### Cell rupture

the tissue maceration was the first step in the DNA extraction from plant cells. The liquid nitrogen/extraction buffer are responsible for breaking the membranes and releasing the cellular content (DNA, proteins, etc).

✓ Three hundred milligram of plant tissue (i.e., matured and young leave, roots, stems and seeds) from each was grinded in liquid nitrogen with pestle and mortal and the grinded sample was divided in to three and each transferred to 2 ml tubes. Seeds were grinded twice just before and after addition of liquid nitrogen. This is to facilitate the breaking down of the endosperm.
✓ 800μl of CTAB buffer solution was added in each tube and incubate for 1 hour at 65 °C in a recirculating water bath with gentle shaking in every 10 minute.
✓ After the incubation period, 400μl of chloroform: isoamyl (ratio 24:1) was added to the tube again and mixed for about 50-70 times; finally the tubes were centrifuged at 13000 rpm for 10 minutes. Then the supernatants were transferred to a new 1.5 ml tubes.

#### DNA precipitation and purification

The DNA pellet was washed to purify to the maximum.

✓ 50μl of 7.5M Ammonium Acetate followed by 500μl ice cold absolute ethanol was added to the recovered supernatant. The samples in the tubes were manually homogenize and placed in the freezer at −20°C for 1 hour. Then the tubes were centrifuge for 5 minutes at 13000 rpm. At this point the DNA pellet remain attached to the tube wall in the bottom, and the liquid was carefully disposed.
✓ To improve the washing of the pellet, 500μL of 70% ethanol was added and manually and gently agitated until the pellet peels off the wall. Then the tubes were centrifuged at 13000 rpm for 1 minutes and the alcohol disposed. Washing was done twice with the same procedure.

#### DNA elution

at this stage the DNA pellet was dried and re-suspended in aqueous solution.

✓ After disposal of all the liquids the tubes were open at room temperature for air drying the DNA for overnight. Once the pellet is completely dry and free from ethanol, the pellets were dissolved in 30μl of nuclease free water.
✓ Finally the tubes were stored in a freezer (−20 °C) for preservation.

### 3.3. Evaluation of DNA extraction

#### 3.3.1. Quantification and integrity of DNA

Quantity or concentration of DNA was determined by Qubit as described by (Zonta *et al*., 2015). Moreover, the size and integrity of the genomic DNA were confirmed by agarose gel electrophoresis. It was analyzed by agarose gel electrophoresis using 0.7% agaros for genomic DNA. Electrophoresis was performed using 1× Tris–Acetate EDTA (TAE) buffer and a constant voltage of 100 V for 40min. The DNA bands were visualized and images were acquired using UV trans-illuminator. Moreover, statistical analysis was carried out using gen-stat software.

#### 3.3.2. PCR conditions

PCR-based amplification of DNA was carried out in a 20 litre reaction mixture. The reaction mixture contained the following: 4μl 5xFIREPol master mix (FIREPol DNA polymerase, 5x Reaction buffer, 12.5 mMMgCl2 and 1mMdNTPs of each), 0.5μl forward and reverse primer each, 14μl H_2_O and 1μl template DNA). Amplification of the DNA was performed in a thermo-cycler in the following manner: initial denaturation at 95°C for 5 min, followed by 35 cycles of denaturation at 95°C for 1 minute, primer annealing at 58°C for 1 min, and extension at 72°C for 2 min, with a final extension at 72°C for 10 min. The reaction mixture was stored at 4°C, until it was loaded onto the gel. The PCR products were loaded using loading dye on 2.96% agarose gel mixed with Edithium bromide in 1X TAE buffer, and were visualized under UV light. Similar to that of the genomic DNA, the gel was photographed using the UV gel documentation system.

## 4. Result and Discussion

### 4.1. Genomic DNA yield

The result of this study showed that DNA was successfully extracted from all tissues of plants which is in concordance with Tamari *et al*. (2013) and the yield of DNA obtained was variable ranging from 179 ng μl^-1^ in stem to 750 ng μl^-1^ in young leave (Table 1). It is within the range of 175 ng μl^-1^ to 1075 ng μl^-1^ as reported by Tamari *et al*. (2013) based on their study on Petunia hybrida tissues. But Sahu *et al*. (2012) reported DNA concentration ranged from 880 to 990 ng μl^-1^ in mangroves plants which is beyond the range of this research finding. This indicates that plant tissues were the most important source of variation for DNA yield and can be explained by either some intrinsic tissue-specific variation (e.g., number and size of cells; ratio of mitotic to interphase nuclei; and amount of extra nuclear DNA) or by a particular differences in the structure or biochemical composition (polysaccharides, polyphenols and other secondary metabolites) of the tissue as described by Aboul-Maaty and Oraby (2019).

**Table 1.** Comparison of mean concentrations of DNA (ng/µl) isolated from different tissues of barley crop Using CTAB method

| Tissues | N | $\bar{x}$ ng $\mu\text{l}^{-1}$ | ± SD |
| --- | --- | --- | --- |
| Young leave | 3 | 750a | 121.2 |
| Matured leave | 3 | 710a | 52.92 |
| Seed | 3 | 238b | 218.4 |
| Root | 3 | 206b | 138.3 |
| Stem | 3 | 179b | 83.72 |

*Letter showed significance difference among tissues using Duncun; 'n' number observations, $\bar{x}$ mean, ng $\mu\text{l}^{-1}$ concentration in nano gram per micro litter, 'SD' standard deviation*

DNA yield (Table 1) from young and matured leave was higher than from the roots, seeds and stems. This suggested that this DNA extraction protocol is effective to isolated DNA from matured and young leave. Similar result was also described by Marsal *et al*. (2013) based on their studies on woody plants. Similarly, Marsal *et al*. (2011) reported that young and matured leave provide good result than the recalcitrant tissues (seeds and stems). Steenkamp *et al* (1994) indicated that, young leave are best tissue for good quantity DNA extraction from grape. Tamari *et al*. (2013) reported that DNA isolated from matured and apical leave have good concentration than other tissues ranging from 250-525 and 650-1125 ng μl^-1^, respectively. This is because leave possess more actively dividing cells than other tissues and therefore, have a higher DNA content or cell (Ulloa, 2020). Unlike to this study, Boiteux *et al*. (1999) reported that better quantity of DNA is obtained from the seed of carrot; the authors also indicated that DNA obtained from roots has low yield which is similar to this study. Elsalam *et al*. (2011) reported that 100-200ng of DNA was produced per 100gm of wheat seed sample which is almost in line with this research finding. It is difficult to isolate good quantity of DNA from plant tissues where there are high contaminants like secondary metabolites, proteins, polyphenols and polysaccharides which are mainly concentrated in roots and seeds (Friar, 2005; Varma *et al*., 2007) and due to lignified tissues in stem (Lewinsohn *et al*., 1994).

### 4.2. Integrity of genomic DNA extracted

Banding pattern was affected by tissue source (Figure 1). Degraded or partially degraded DNA preparations were observed in lane 5 (stem) and lane 13 (seed) tissue while best DNA integrity was observed from young and matured leave and this is identical with the result of Marsal *et al*. (2013). The DNA obtained from roots and stems showed very faint bands upon gel electrophoresis probably due to the low concentration of DNA which could be because of the lignified tissue and the presence of higher levels of secondary metabolites (Hasan *et al*., 2012; Varma *et al*., 2007). Moreover, highly viscous and sticky with brownish pellet was observed in the roots and seeds after extraction which is an indication of contamination by phenolic compounds mentioned by Moreira and Oliveira (2011). The pellets from stem, matured and young leave were white with no visible coloration indicating no phenolic contaminants as reported by Sahu *et al*. (2012). Indeed there are report by Moreira and Oliveira (2011), matured leave can produce brownish DNA pellets which is in contrary with this research finding. In general as mentioned by Satyanarayana *et al*. (2017) the clear band in gel depicts the genomic DNA that is sufficiently pure and unsheared for further molecular studies. With this regard DNA obtained from young and matured leave are sufficient enough for PCR amplification.

**Figure 1.**
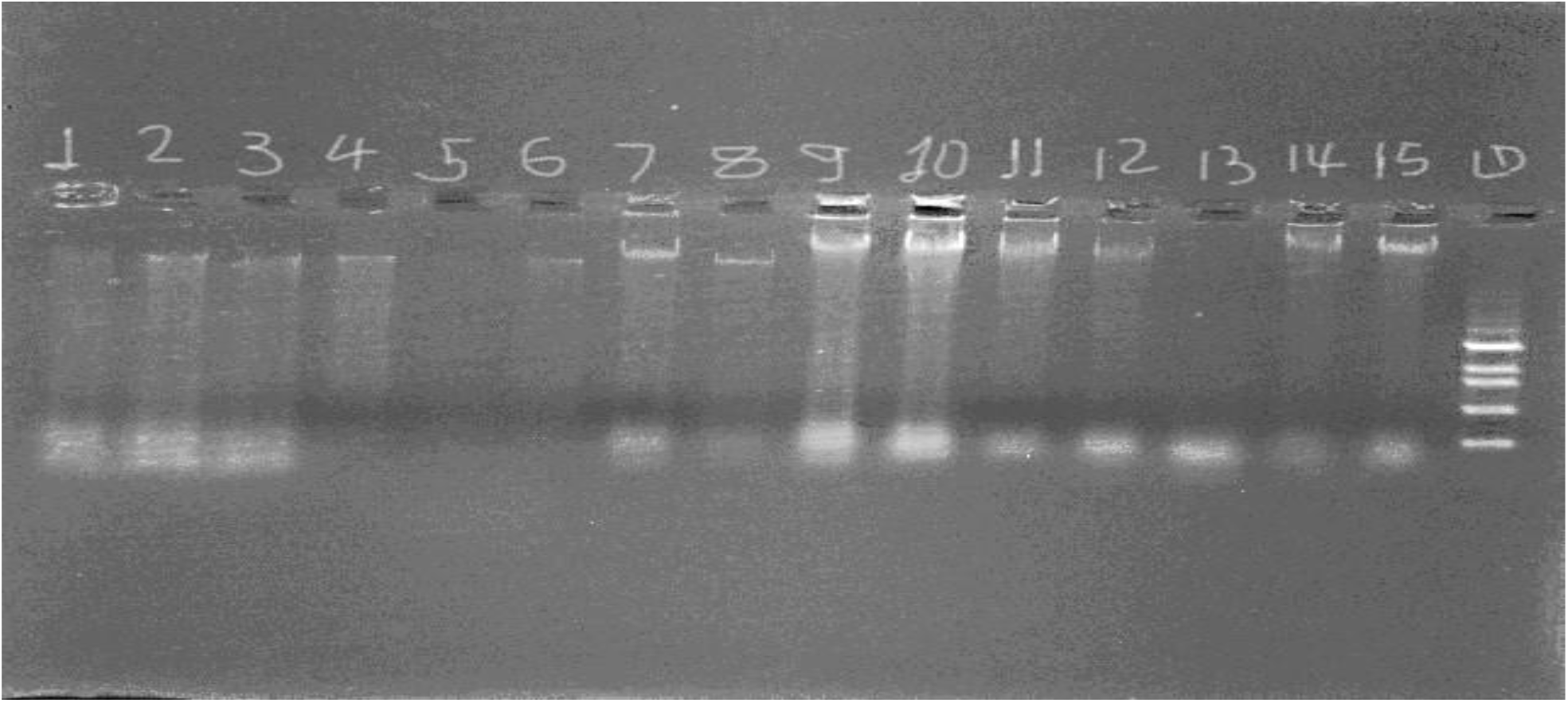
Agarose gel-electrophoresis of barley DNA extracted from its root, stem, old leaves, young leaves and seeds. Lanes 1-3 represent root: lanes 4-6 stem; lanes 7-9 old leaves; lanes 10-12 young leaves, lanes 13-15 seeds and the last is Marker.

### 4.3. Influence of tissue type and developmental stage on PCR efficiency

In order to confirm the quality of the extracted DNA and reliability of the DNA extraction protocol, DNA of each tissue of barley was amplified using primer “HdAMYB”. As it is described by Mei *et al*. (2012) its allelic length was varied from 199-221bp and responsible to amylase gene (Bmy1) in barley. The relative amount of each PCR product estimated by comparing the PCR product with the 200-bp DNA fragment of the 100-bp ladder.

The DNA extracted from matured and young leave using the prescribed protocol was suitable for PCR and showed high band intensity upon gel electrophoresis after amplification with HdAMYB primers **(Figure 2)**. This indicates that the isolated DNA was free from interfering compounds such as secondary metabolites, proteins, polyphenols and polysaccharides (Angeles *et al*., 2005), and it would be suitable for other down streaming processes. Similar research by Moreira and Oliveira (2011) indicated that PCR performed with DNA obtained from young leave of *D. mollis* was successful and produced strong bands for RAPD markers. This indicates that the DNA obtained from young leave was pure enough to PCR amplifications. Most research indicates that DNA obtained from matured leave has poor amplification performance (Williams and Ronakd., 1994; Sahu *et al*., 2012; Varma *et al*., 2007; Elger *et al*., 2009; Moreira and Oliveira, 2011; Sahu *et al*., 2012). However, based on this research it can be possible to obtain quality DNA from matured leave revealing that the DNA extraction method was efficient enough to get quality DNA from matured leave that can be amplified through PCR.

**Figure 2.**
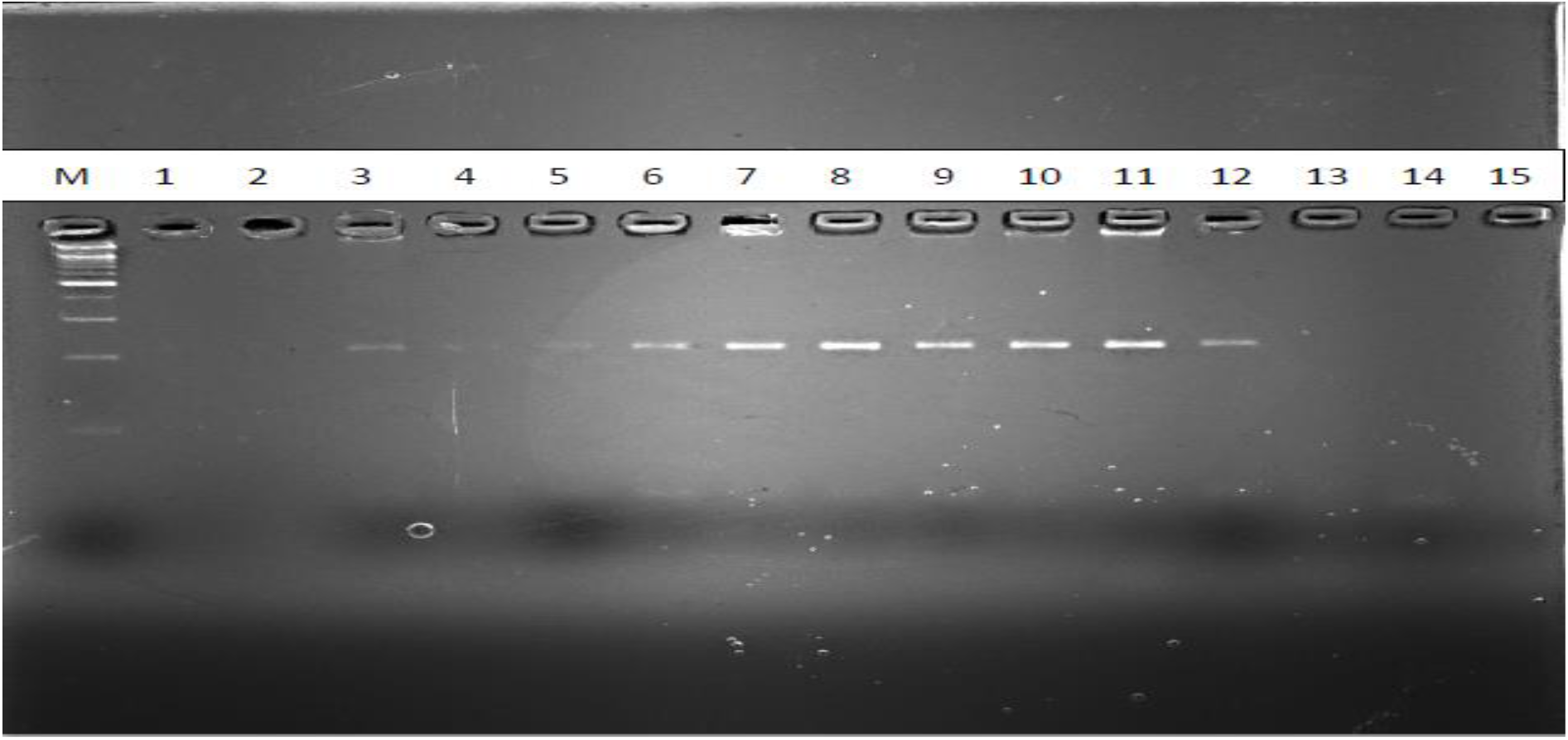
PCR Amplified DNA visualized by gel electrophoresis. M marker (ladder) Lanes 1-3 represent root; lanes 4-6 stem; lanes 7-9 old leaves; lanes 10-12 young leaves, lanes 13-15 seeds

Amplification was also possible with DNA obtained from stem though it was very faint probably due to low concentration of DNA and the lignified tissue of the stem (Lewinsohn *et al*., 1994; Schori *et al*., 2013). In contrast to this, Aydın *et al*. (2020) reported that, stem can produce good quality DNA for PCR amplification as equally as to that of leave. Lucas *et al*. (2019) indicated that, DNA obtained from stem can be used as alternative source of DNA and can be used for loci amplification in palm trees. On the other hand, there was unreliable amplification from the root source while there was no amplification from the seed source. This may be due to the fact that roots are more susceptible to fungal contamination than other plant parts (Di Bonito *et al*., 1995) and can be exposed to humic acids from soil (Puglisi *et al*., 2013). Boiteux *et al*. (1999) also mentioned that poor amplification was observed with DNA isolated from carrot roots. Regarding to the seeds, Boiteux *et al*. (1999) reported that except standard CTAB (cetyltrimethylammonium bromide) buffer together with organic solvents all other methods could not isolate DNA from the seed that can be amplified. Additionally, Hasan *et al*. (2012) extracted high quality DNA from the seed using modified CTAB. However, the CTAB protocol described in this research and adopted by AASTU failed to produce quality DNA from seeds which could be used for PCR amplification.

## 5. Conclusion

Plant tissues were found an important source of variation for DNA isolation and this research tried to describe the best tissues for isolation of DNA from barley crop. The best quality and quantity of DNA was isolated from matured and young leave of barley than other tissues (seed, stem and roots) with the modified protocol. Though the concentration and the band intensity of genomic DNA is weak, PCR amplification was possible with the DNA obtained from the stem and hence can be used as alternative source of DNA where there is lack of access for the leaves. But it is impossible to use roots and seeds as source of DNA with the method prescribed here in the research. Hence, it may need another optimization or modification of the protocol for good quantity and quality DNA extraction from the roots and seeds suitable for PCR amplification.

## Acknowledgment

I would like to acknowledge Mrs. Meriam, Mr. Ashebir and Mr. Minilu for their valuable advice, guidance and material support during the course of the laboratory activities. It is also necessary to extend the acknowledgement for Debre birehan Agricultural research Centre for sourcing the barley seed. Finally, I am very much indebted to thank the department of biotechnology itself for having all the laboratory facilities and allowing me to do this research.

## Authors’ contributions

Nigussie K. wrote the whole manuscript and Gizachew H. and Hemaltha P. did the review and editing process.

## Annexes

Annex 1: Materials and Reagents used

|  |  |
| --- | --- |
| Microfuge | Micropipetor |
| Microfuge tube | Electric power source |
| Mortar and pestle | Parafilm |
| Agarose gel electrophoresis system | Thermal cycler |
| Erlenmeyer flask, | Eppendorf (PCR) tube |
| Microwave oven, | Qubit tube |
| 65°C water bath | Liquid nitrogen |
| CTAB buffer | Absolute ethanol (ice cold) |
| 70% Ethanol (ice cold) | Taq polymerase |
| 7.5M ammonium Acetate | dNTPs |
| Tris Acetate EDTA buffer | MgCl<sub>2</sub> |
| DNA size marker | Sterile water |
| Chloroform; Iso Amyl Alcohol (24:1) | Agarose 1x loading buffer |
| Primers | Gel red |

Annex 2: Preparation of working solution

CTAB preparation and its components

➣ 2gm CTAB (hexadecyltrimethyl-ammonium bromide
➣ 10ml of 1M Tris pH 8.0
➣ 4ml of 0.5 EDTA pH 8.0 (Ethylene diamine tera Acetic acid)
➣ 28ml of 5M NaCl
➣ 40ml H_2_O
➣ 1g PVP 40 (polyvinyl pyrrolidone vinylpyrrolidinehomopolymer) Mw 40000) adjusted all to pH 5.0 with HCl and make up to 100ml with H_2_O 1M Tris pH 8.0 preparation
➣ 121.14 gm of Tris base was dissolved in 50ml H_2_O and adjusted to pH 8.0 by adding concentrated HCl Chloroform isomyl alcohol preparation
➣ 96 ml of chloroform was mixed with 4ml of isoamyl alcohol in the ratio of 24:1 Preparation of 7.5M of Ammonium Acetate
➣ 28.905gm of Ammonium acetate was dissolved in 50ml of H_2_O 50x TAE buffer preparation
➣ 242g Tris was dissolved in 500ml H2O
➣ 100ml of 0.5M EDTA (pH 8.0) was added
➣ 57.1 ml of glacial acetic acid was also added
➣ And finally the total volume was adjusted to 1 liter with distilled water 1 liter 1x TAE buffer preparation
➣ 20 ml of 50x TAE buffer was added to 980ml of distilled water

